# Probing the activity of cysteine cathepsins in inflammatory bowel diseases

**DOI:** 10.1101/2025.09.18.677249

**Authors:** Bethany M. Anderson, Alexander R. Ziegler, Rhiannon I. Campden, Hongyi Wu, Bangyan Xu, Rachel M. McQuade, Simona E. Carbone, Daniel P. Poole, Alan E. Lomax, David Reed, Stephen J. Vanner, Robin M. Yates, Nigel W. Bunnett, Laura E. Edgington-Mitchell

## Abstract

Cathepsin S is a cysteine protease that has been implicated in inflammatory bowel diseases (IBD) for its ability to promote visceral pain. Given its pro-inflammatory roles, we hypothesized that cathepsin S would drive other symptoms associated with IBD. Using activity-based probes, we investigated cysteine cathepsin activation in human and murine colitis. We observed a significant increase in fecal cathepsin S in patients with ulcerative colitis compared to healthy controls, while cathepsin S in mucosal biopsies was unchanged. Mice with experimental colitis exhibited a modest increase in mucosal activity of both cathepsin S and X compared to naïve mice. Luminal secretion of cathepsin S was dramatically increased upon colitis induction, although differences between mouse colonies were observed. To investigate the contribution of cathepsin S and cathepsin X to colitis, we induced colitis in cathepsin-deficient mice. Cathepsin X-deficient mice exhibited no clear differences in disease indicators compared to wild-type mice. While cathepsin S-deficient mice exhibited less rectal bleeding, less splenomegaly and marginally improved histological scores, weight loss, diarrhea, colon shortening, and myeloperoxidase activity were not significantly different from wild-type mice. To determine whether pharmacologic inhibition of cathepsin S activity would ameliorate symptoms of colitis, a reversible inhibitor LY3000328 was administered to mice at the initiation of colitis. LY3000328 provoked a clear upregulation of cathepsin S and L activity in the mucosa, most likely through a compensatory mechanism. This increase in protease activity was associated with exacerbated histological scores and splenomegaly. Collectively, these results suggest that cathepsin S, but not cathepsin X, may contribute to some of the symptoms of experimental colitis. While cathepsin S has potential to be a therapeutic target in colitis, improved strategies to sustain its inhibition are required in future.

## Introduction

Inflammatory bowel diseases (IBD), including ulcerative colitis (UC) and Crohn’s disease (CD), are characterized by relapsing and remitting bouts of diarrhea, rectal bleeding, increased urgency and pain. The etiology of IBDs is not well understood. It has been hypothesized that damage to the intestinal epithelium leads to impaired barrier function, leakage of microbes into the mucosa and activation of pro-inflammatory neutrophils, macrophages, and T lymphocytes. Macrophages are a rich source of cysteine cathepsin proteases, which contribute to inflammation associated with diverse pathologies^1^.

Cathepsin S is a lysosomal cysteine protease associated with inflammation. Among its substrates are protease-activated receptor 2 (PAR_2_), where cleavage activates the receptor in a biased manner^2,3^, and invariant chain, which promotes MHC II maturation and antigen presentation^4^. Compared to other cysteine cathepsins, cathepsin S has a relatively broad pH range and is able to maintain its enzymatic activity following release from cells into the extracellular milieu^5^.

Using a model of piroxicam-induced colitis in IL-10-deficient mice, cathepsin S was previously demonstrated to be activated in the proximal colon, cecum, and luminal fluids^6^. Cleavage of PAR_2_ by cathepsin S results in hyperexcitability of nociceptive dorsal root ganglia neurons that innervate the colon. Colonic infusion of cathepsin S increased the visceromotor response to colorectal distension in wild-type mice, which indicates heightened sensitivity to innocuous and painful stimuli. This response was attenuated in mice lacking PAR_2_. Cathepsin S-deficient mice with trinitrobenzenesulfonate (TNBS)-induced colitis exhibited reduced visceromotor response to colorectal distension compared to wild-type mice. Collectively, these results indicate a role for cathepsin S in provoking inflammatory pain associated with colitis.

In addition to visceral pain, activation of PAR_2_ is known to provoke other features of colitis, including dysmotility and secretion, loss of barrier function, submucosal edema, cytokine production, and inflammatory infiltration^7–10^. Accordingly, antagonism of PAR_2_ or inhibition of other PAR_2_-activiting proteases has resulted in improved outcomes in rodent models of experimental colitis^11,12^. Whether cathepsin S contributes to the development of colitis and associated symptoms through its actions on PAR_2_ or other targets has not been investigated in detail.

Steimle and colleagues demonstrated that negative regulation of cathepsin S activity by commensal bacteria such as *Bacteroides vulgatus* may be important for maintaining intestinal equilibrium^13^. In the presence of symbiotic bacteria, cathepsin S may be inhibited by endogenous inhibitors such as cystatin C. Reactive oxygen species induced by a bacterial pathobionts such as *E. coli* mpk provoke homodimerization of cystatin C, rendering it unable to bind to cathepsin S. Elevated cathepsin S activity in the presence of pathogenic bacteria may therefore contribute to intestinal host inflammatory responses. In CD4^+^ T cell-mediated colitis in Rag1-deficient mice, administration of *B. vulgatus* or a cathepsin S inhibitor LY3000328 had similar effects in attenuating weight loss, improving histological scores, and reducing numbers of CD3^+^CD4^+^ T cells and MHC-II^hi^CD11c^+^ dendritic cells, providing additional support for this hypothesis.

To investigate the contribution of cysteine cathepsins to colitis in more detail, we applied activity-based probes to measure their proteolytic activity in an acute model of experimental colitis induced by dextran sulfate sodium (DSS). We then investigated the effects of cathepsin deletion and on colitis development and associated symptoms.

## Experimental Procedures

### Human samples

Written and verbal consent were obtained prior to enrolment, and all protocols were approved by and carried out in accordance with the Queen’s University Human Ethics Committee. Patients were well-characterized individuals with active UC or CD (refer to Table 1-2). Healthy individuals were volunteers experiencing no gastrointestinal abnormalities or abdominal pain. For UC and CD patients, biopsies were obtained from sites of active inflammation. Fresh biopsies were washed in PBS and then snap frozen for protease analysis. Fecal samples were homogenized in 0.7% NaCl (0.5 g per 4 mL) and centrifuged to clear solids. Supernatants were frozen for protease analysis.

**Table 1.**
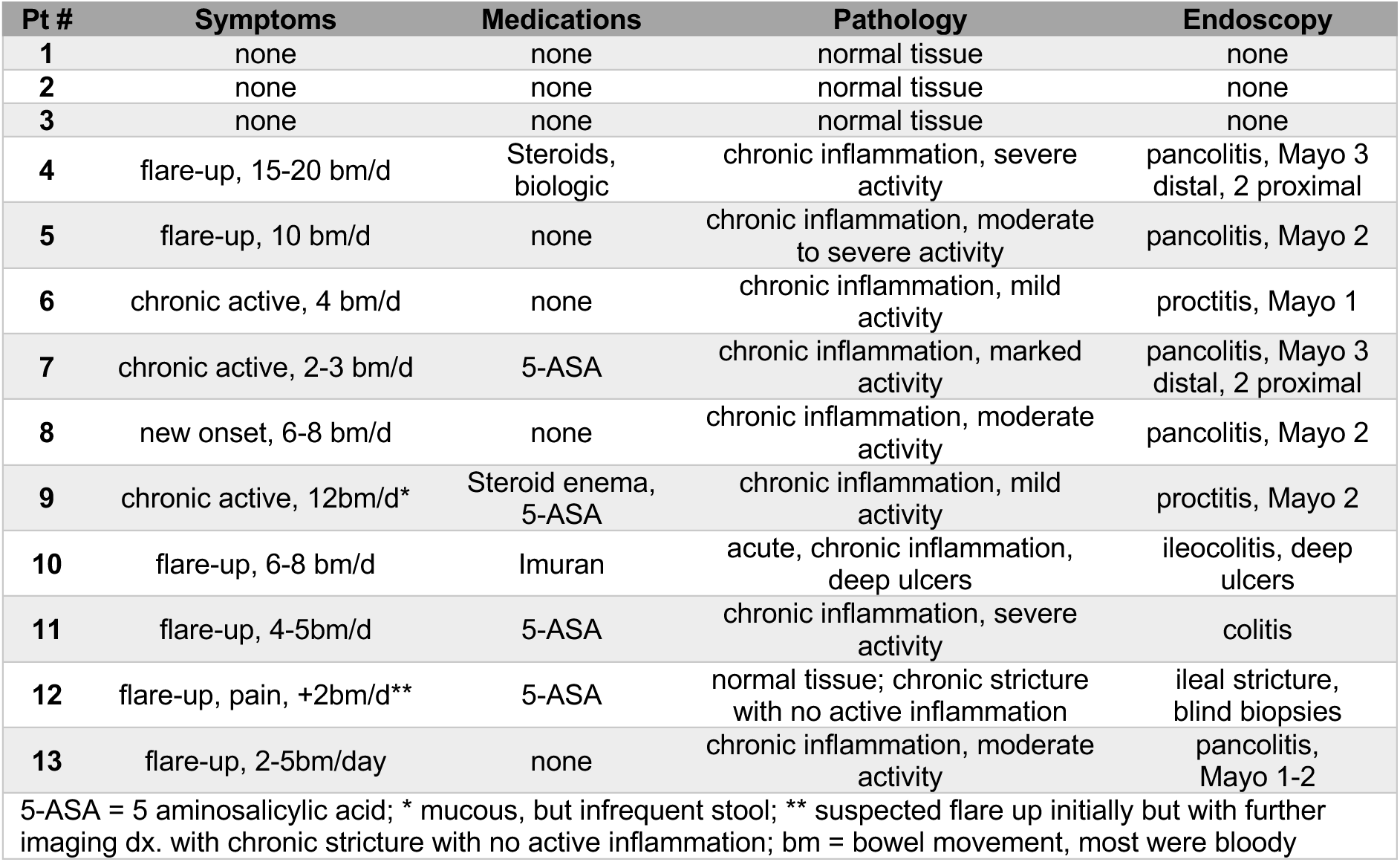
Profiles of patients from which mucosal biopsies were collected.

| Pt # | Symptoms | Medications | Pathology | Endoscopy |
| --- | --- | --- | --- | --- |
| 1 | none | none | normal tissue | none |
| 2 | none | none | normal tissue | none |
| 3 | none | none | normal tissue | none |
| 4 | flare-up, 15-20 bm/d | Steroids, biologic | chronic inflammation, severe activity | pancolitis, Mayo 3 distal, 2 proximal |
| 5 | flare-up, 10 bm/d | none | chronic inflammation, moderate to severe activity | pancolitis, Mayo 2 |
| 6 | chronic active, 4 bm/d | none | chronic inflammation, mild activity | proctitis, Mayo 1 |
| 7 | chronic active, 2-3 bm/d | 5-ASA | chronic inflammation, marked activity | pancolitis, Mayo 3 distal, 2 proximal |
| 8 | new onset, 6-8 bm/d | none | chronic inflammation, moderate activity | pancolitis, Mayo 2 |
| 9 | chronic active, 12bm/d* | Steroid enema, 5-ASA | chronic inflammation, mild activity | proctitis, Mayo 2 |
| 10 | flare-up, 6-8 bm/d | Imuran | acute, chronic inflammation, deep ulcers | ileocolitis, deep ulcers |
| 11 | flare-up, 4-5bm/d | 5-ASA | chronic inflammation, severe activity | colitis |
| 12 | flare-up, pain, +2bm/d** | 5-ASA | normal tissue; chronic stricture with no active inflammation | ileal stricture, blind biopsies |
| 13 | flare-up, 2-5bm/day | none | chronic inflammation, moderate activity | pancolitis, Mayo 1-2 |

5-ASA = 5 aminosalicylic acid; \* mucous, but infrequent stool; \*\* suspected flare up initially but with further imaging dx. with chronic stricture with no active inflammation; bm = bowel movement, most were bloody

### DSS-induced colitis

All animal studies were approved by and carried out in accordance with the guidelines of the Animal Ethics Committee at Monash University or the Animal Care and Use Committee at the University of Calgary. Wild-type C57Bl/6J mice were purchased from the Monash University in-house colony. Wild-type and Cathepsin S-and cathepsin X-deficient C57Bl/6J mice, WT, *Ctss^-/-^* and *Ctsz^-/-^*respectively, were bred at the University of Calgary^14,15^. All mice were maintained with free access to food and water under temperature and light controlled conditions. Colitis was induced in 8-10-week-old male mice by administering dextran sulfate sodium (MP Biomedicals 36,000-50,000 Da; 3% unless otherwise indicated) in the drinking water for 6 days. Control mice received drinking water alone. Body weights and symptoms were monitored daily. Fecal pellets were scored using scales for consistency (0 – dry; 1 – moist but firm; 2 – soft but still formed; 3 – unformed but bulky; 4 – liquid) and blood (0-no blood; 1 – subtle blood or darkening of the stool; 2 – red streaks in stool; 3 – obvious rectal bleeding). Fecal samples were collected and frozen daily. Unless otherwise indicated, mice were humanely killed on day 6. Colons were harvested and flushed with phosphate buffered saline (PBS). Luminal fluid was collected, centrifuged to remove solids, and frozen. Proximal colon tissue was divided and frozen for gel analysis and myeloperoxidase (MPO) assay or fixed for microscopy or histology. Spleens were also harvested and frozen. For the cathepsin S inhibitor trial, 2% DSS was used. LY3000328^16^ (30 mg/kg in 200 µl 85% NaCl (0.9%), 10% DMSO and 5% Tween-20) or vehicle control was administered daily by intraperitoneal injection from the introduction of DSS.

### Ex vivo tissue imaging

On day 6, naïve or DSS-treated mice were injected with BMV109^17,18^ or BMV157^19^ intravenously by tail vein (20 nmol in 100 µl 20% DMSO/PBS). After six hours, colons were harvested, flushed with PBS, and imaged for Cy5 fluorescence using an IVIS Lumina XR III in vivo imaging system (Perkin Elmer). Proximal colons were cut along the mesenteric border and pinned onto Sylgard-lined 35 mm dishes (mucosa down) for fixation in 4% paraformaldehyde overnight at 4°C. Whole mount preparations were washed with PBS azide (0.1%) and blocked for 1 hour in 10% normal horse serum and 0.1% Triton X-100 in PBS azide. Preparations were washed 3 times with PBS azide and then incubated overnight at 4°C with rat anti-CD68 (1:500; clone FA-11; BioLegend; 137001) in blocking buffer. After a further 3 washes, donkey anti-rat-Alexa Fluor488 (1:500; Life Technologies; A-21208) in PBS azide was applied for 1 h at RT. After 3 additional washes, whole mount preparations were mounted using ProLong Gold (Invitrogen) and imaged with a Leica SP8 confocal microscope.

### Analysis of cathepsin probe labeling by SDS-PAGE

Mouse proximal colon tissues or human mucosal biopsies were lysed by sonication in PBS (for in vivo-labeled samples) or citrate buffer (for in vitro-labeled samples; 50 mM citrate, pH 5.5, 0.5% CHAPS, 0.1% Triton X-100, 4 mM DTT)^18,20,21^. Lysates were cleared by centrifugation at 14,000 x g for 5 min at 4°C. Cleared luminal fluids were concentrated using a 3-KDa cutoff column (Amicon). Fecal samples were homogenized in PBS and centrifuged to clear solids. Total protein concentration was measured in colon lysate, luminal fluid and fecal supernatant using a BCA assay (Pierce) and diluted into PBS or citrate buffer (50 µg total protein in 20µl). For in vivo-labeled samples, 5x sample buffer was immediately added (50% glycerol, 250 mM Tris-Cl, pH 6.8, 10% SDS, 0.04% bromophenol blue, 6.25% β-mercaptoethanol, diluted to 2x). For in vitro-labeled lysates or fecal supernatants, BMV109 (0.5 µM) or BMV157 (1 µM) were added from a 100x DMSO stock concentration and incubated at 37°C for 30 min. Where indicated, LY3000328 (10 nM – 100 µM) or MDV590^22^ (10 µM) was added 30 min prior to probe addition. Labeling reaction was quenched by addition of 5x sample buffer. Samples were boiled for 5 min and resolved on a 15% SDS-PAGE gel. Gels were scanned for Cy5 fluorescence using a Typhoon 5 flatbed laser scanner (GE Healthcare). After transferring to nitrocellulose membranes, membranes were immunoblotted overnight with goat anti-cathepsin S (1:1000; R&D Systems; AF1183) or goat anti-cathepsin X (1:1,000; R&D Systems; AF1033) followed by detection with donkey anti-goat-IR800 (1:10,000; LI-COR; 9263-2214). Membranes were scanned using a Typhoon 5 or Odyssey Imaging System (LI-COR).

### Immunoprecipitation

Probe-labeled samples were divided into input and pulldown, each containing 50 µg total protein. Pulldown samples were diluted in 500 µl immunoprecipitation (IP) buffer (PBS, pH 7.4, 0.5% NP-40, 1 mM EDTA) followed by 10 µl of the indicated antibody (all from R&D Systems): anti-cathepsin S (AF1183), anti-cathepsin X (AF1033), anti-cathepsin L (AF1515) or anti-cathepsin B (AF965). These antibodies have been previously well characterized^17,23–26^. Protein A/G agarose beads (40 µl slurry; Santa Cruz Biotechnology) were washed in IP buffer and added. Pulldowns were rotated overnight at 4°C, and supernatants were collected and precipitated in acetone for 1 h at-20°C. Proteins were pelleted and dried prior to dissolving in 20 µl 1x sample buffer. Beads were washed 4 times in IP buffer and once in NaCl (0.9%) and resuspended in 20 µl 2x sample buffer. Input, pulldown and supernatants were boiled for 5 min and resolved by fluorescent SDS-PAGE as above.

### Secretion assay

Proximal colon tissue from naïve and DSS-treated mice were weighed and incubated overnight at 37°C in 500 µl DMEM containing 10% fetal bovine serum. Supernatant was collected, filtered with a 50-kDa cutoff spin column to exclude large serum proteins and concentrated with a 3-kDa spin column (Amicon). Cathepsins were then labeled with BMV109 and analyzed as above.

### Histology

Colon tissues were fixed overnight at 4°C in 4% paraformaldehyde in PBS, paraffin embedded, sectioned and stained with hematoxylin and eosin according to standard protocols. Slides were scanned on a Mirax Digital Slide Scanner (Zeiss). Slides were de-identified and four random regions from each colon were scored for crypt organization (0-5), immune cell infiltration (0-5) and goblet cell expression/cavitation (0-5) based on modified histomorphological evaluation criteria^21,27,28^. All images were assessed while blinded to the assessor.

### Myeloperoxidase activity assay

Tissues from the middle region of the colon were assayed for myeloperoxidase activity. Tissues were sonicated in buffer containing 50 mM potassium phosphate, pH 6.0, 0.5% hexadecyl trimethylammonium bromide (50 mg tissue per mL) and supernatants were cleared by sonication. In a 96 well plate, 7 µl sample was diluted in 193 µl substrate solution containing 50 mM potassium phosphate, pH 6.0, O-dianisidine HCl (0.167 mg/mL) and 0.0005% H_2_O_2_. Absorbance at 460 nm was measured every 30 s for 15 min on a FlexStation 3 (Molecular Devices) and slopes were recorded.

### Statistical analyses

All experiments were performed with at least 3 biological replicates. Data are reported as means ± SEM. Statistical significance was determined by comparing two groups using a Student’s t-test or Mann-Whitney U test, and p values of less than 0.05 were considered significant.

## Results

### Cathepsin S secretion is increased in IBD patients

To determine whether cathepsin S is activated in the colons of patients with ulcerative colitis (**Table 1**), we applied a fluorescently quenched activity-based probe (ABP), BMV157^19^. Cathepsin S binds to BMV157 in an activity-dependent manner, triggering the release of a QSY21 quenching group and subsequent emission of Cy5 fluorescence. Probe binding is irreversible and can be detected by scanning gel-resolved proteins for Cy5 fluorescence or imaging whole tissue or tissue sections. In lysates from mucosal biopsies prepared at pH 5.5 or 7.4, we did not observe any statistically significant differences in BMV157 labeling between healthy volunteers and patients with ulcerative colitis (**Fig 1A-B**). By contrast, in supernatants prepared from fecal samples (**Table 2**), cathepsin S activity was significantly increased in patients with ulcerative colitis (UC) (**Fig 1B,D**), and total cathepsin was increased in both UC and Crohn’s disease samples compared to those from healthy volunteers (**Fig 1C,E**). We confirmed the identity of the BMV157-labeled protease by immunoprecipitating the samples with a cathepsin S-specific antibody (**Fig 1F**). These data suggest that luminal secretion of cathepsin S is increased in patients with active inflammatory bowel diseases.

**Figure 1.**
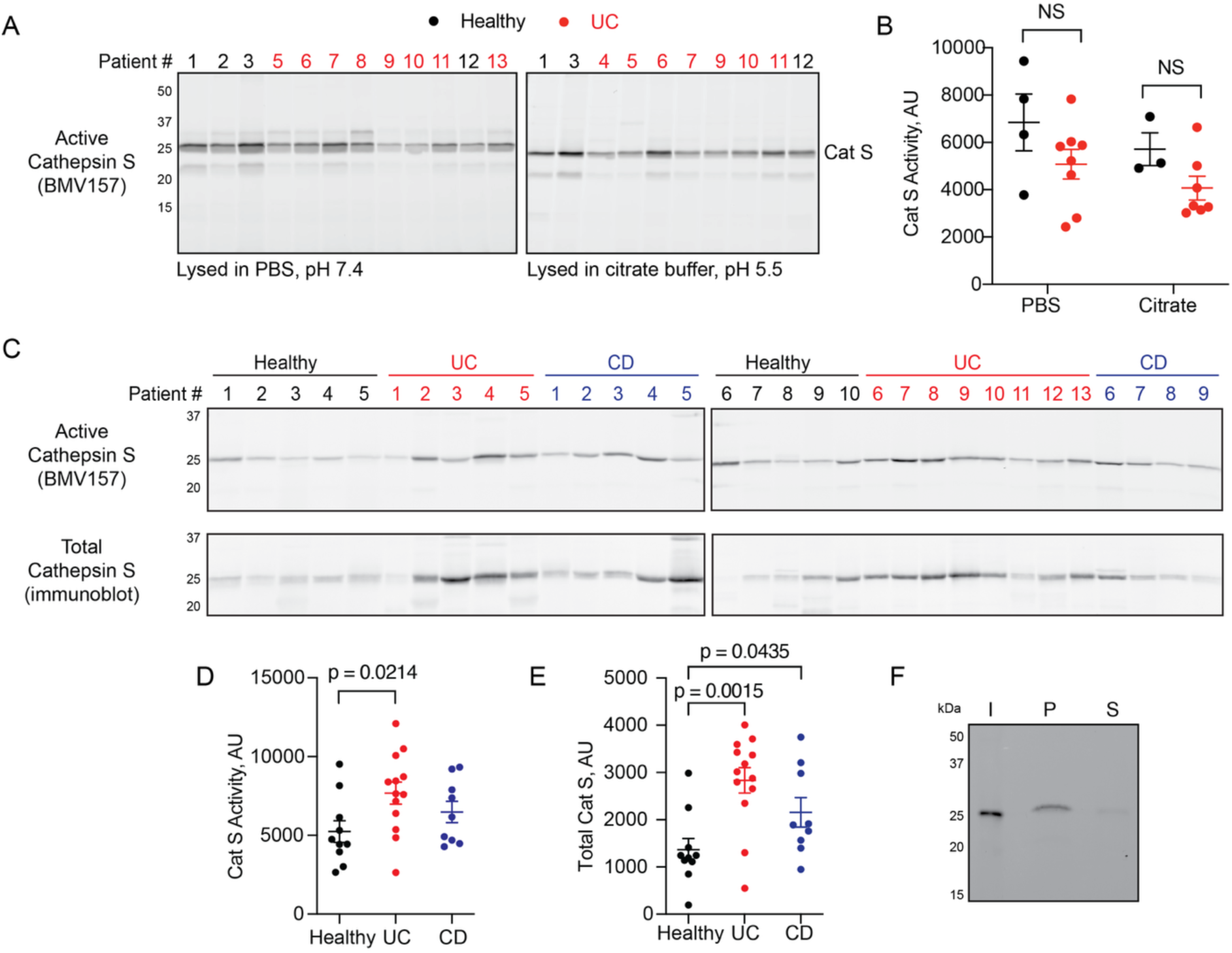
Cathepsin S is increased in fecal samples of patients with inflammatory bowel diseases. **A)** Mucosal biopsies from healthy volunteers or patients with ulcerative colitis (UC) were lysed in PBS, pH 7.4 (left) or citrate buffer, pH 5.5 (right), labeled with BMV157 and analyzed by in-gel fluorescence. **B)** Cathepsin S labeling was quantified by densitometry. Data are represented as means ± SEM (n=3-8; Mann-Whitney U test). **C)** Homogenized fecal samples from healthy volunteers or patients with UC or Crohn’s disease (CD) were labeled with BMV157 followed by analysis of in-gel fluorescence (top) and immunoblotting for cathepsin S (bottom). **D-E)** Densitometry analysis of cathepsin S activity and total levels in C. Data are represented as means ± SEM (n=9-13; Mann-Whitney U test). **F)** Immunoprecipitation of CD sample with a cathepsin S-specific antibody. I = input, P = pulldown, S = supernatant. p values < 0.05 were considered significant.

**Table 2.**
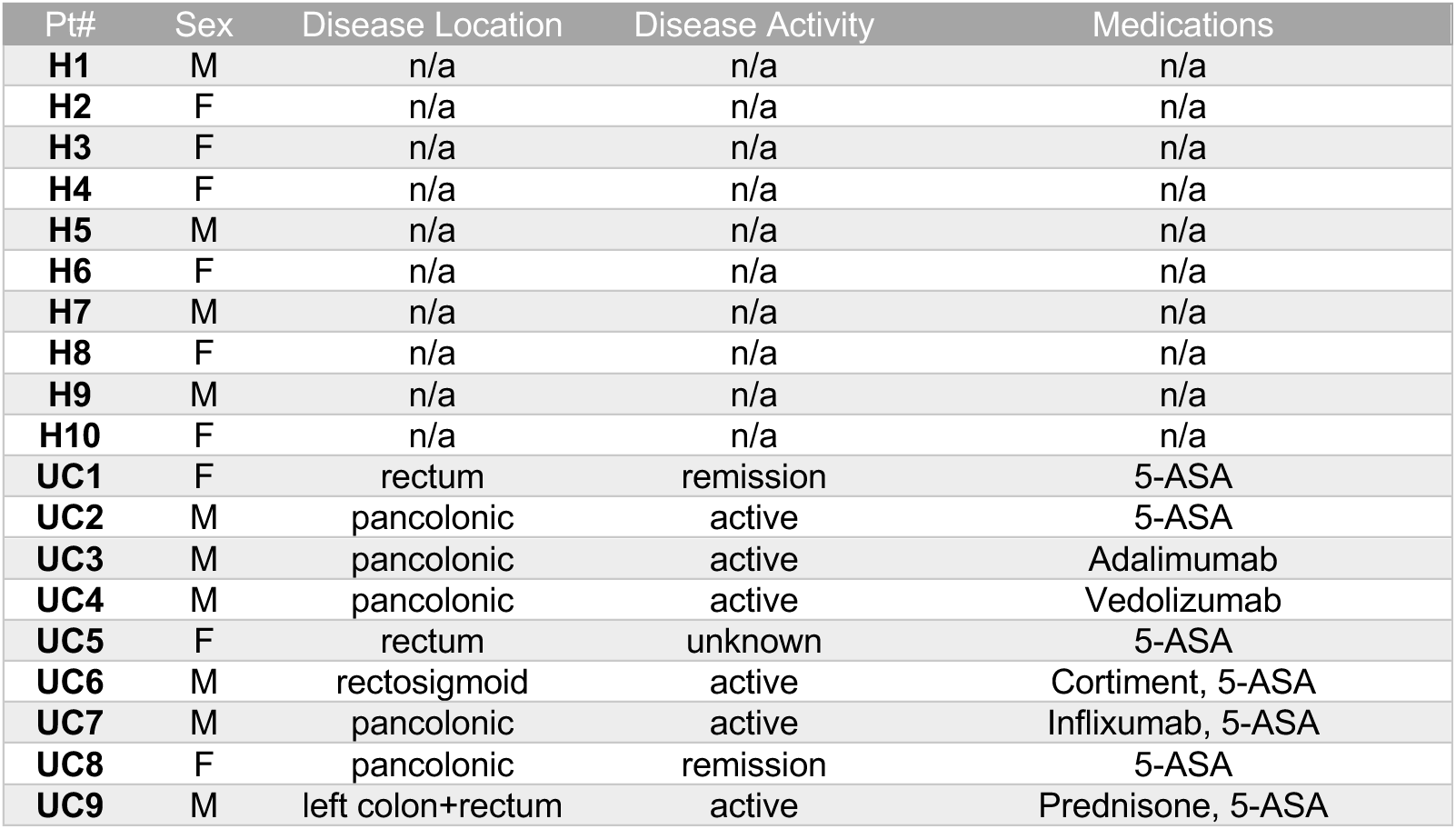
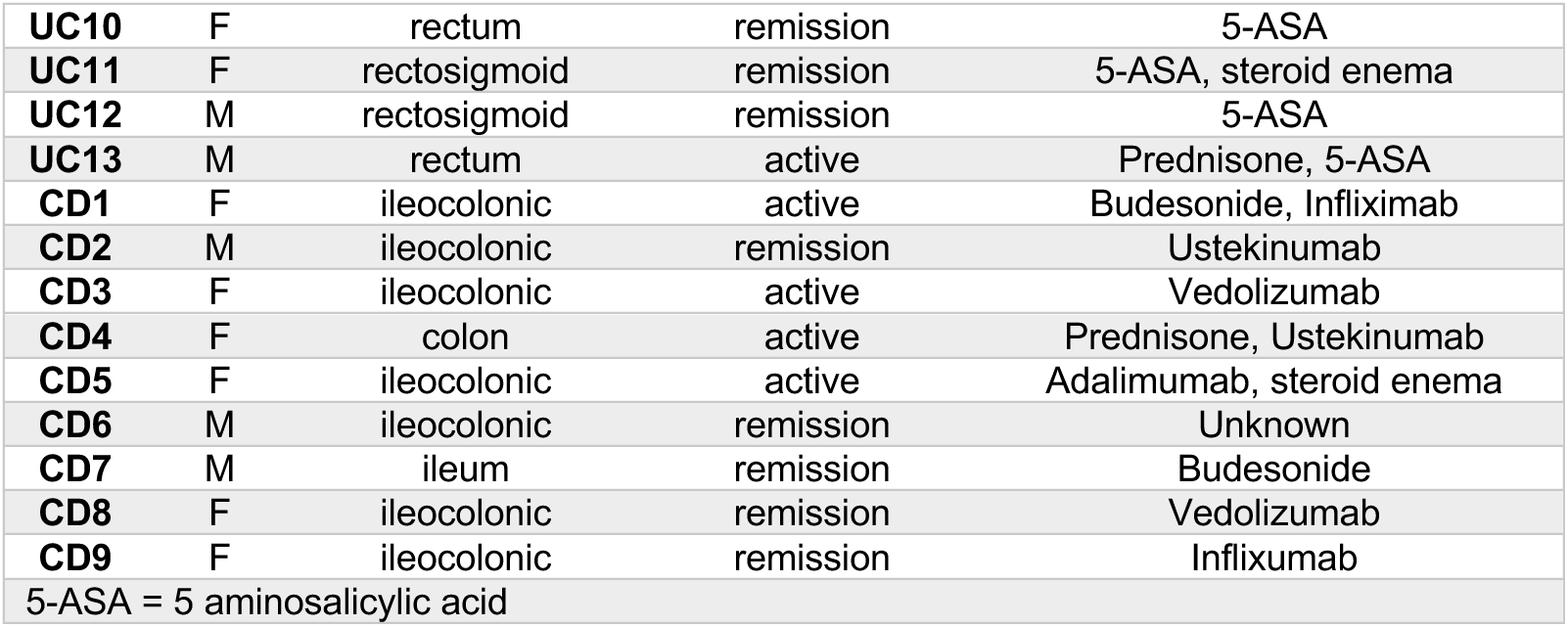
Profiles of patients from which fecal samples were collected.

### Cathepsin S secretion is increased in experimental colitis

We investigated cathepsin S activation in a model of experimental colitis induced by DSS using BMV157 and a pan cysteine-cathepsin ABP BMV109 that binds to cathepsin X, B, S, and L^17^. After in vivo ABP administration, we imaged probe fluorescence in colon tissues ex vivo. With both probes, we observed a significant increase in fluorescence in the proximal region of colons from DSS-treated mice compared to those of naïve mice (**Fig 2A-B**). There was no difference in fluorescence intensity in the distal region of the colons. In whole mount submucosal preparations, most of the cathepsin S-specific signal from BMV157 was confined to CD68^+^ macrophages (**Fig 2C**).

**Figure 2.**
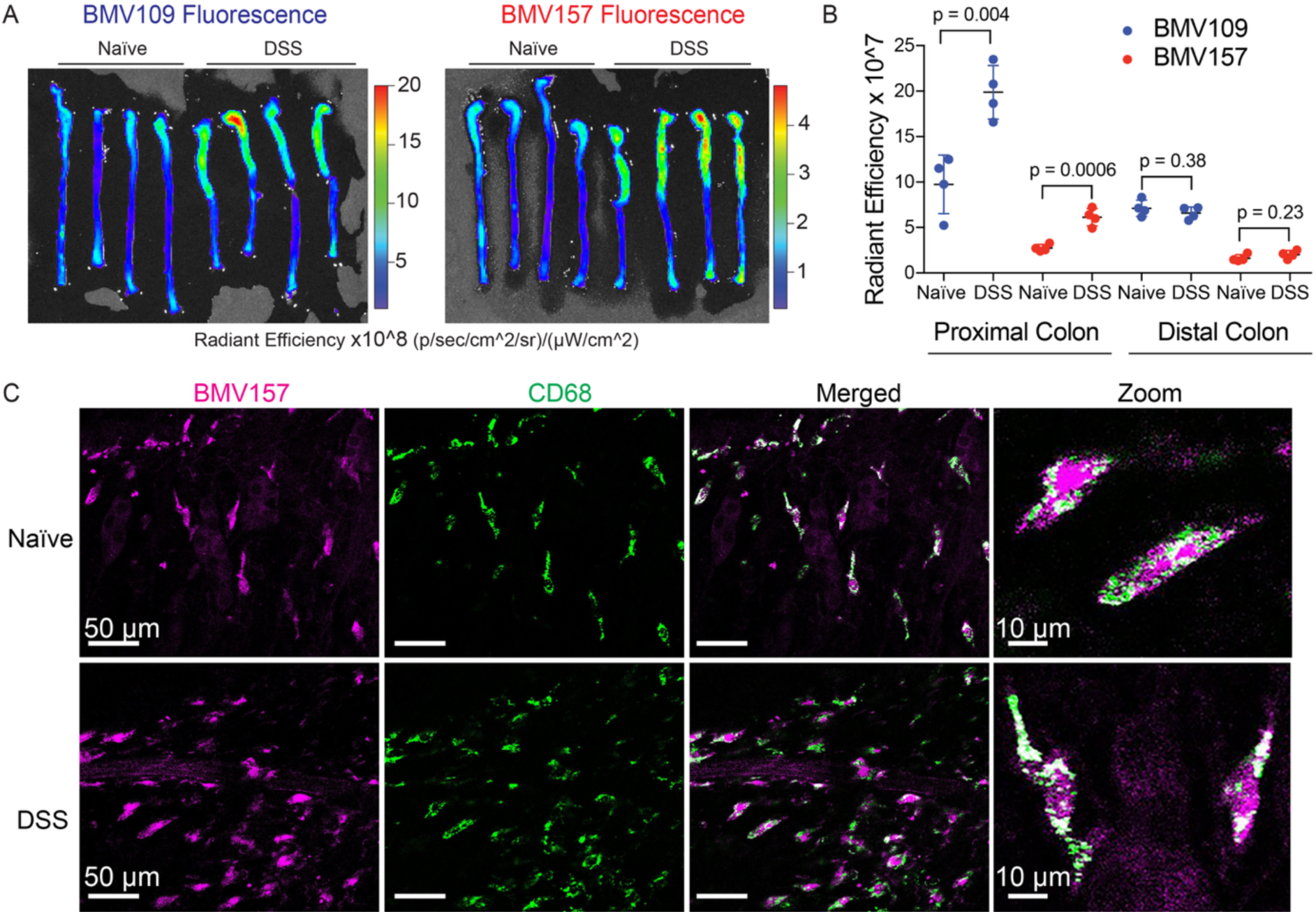
Cathepsin activity is increased in the proximal colon during DSS-induced colitis. **A)** Naïve or DSS-treated mice were injected with BMV109 (pan-cysteine cathepsin probe) or BMV157 (cathepsin S-selective probe) on day 6. Colons were removed and imaged for Cy5 fluorescence. Gains were set for each probe independently. **B)** Quantification of fluorescent signal (radiant efficiency) in proximal and distal colons from (A). Data are represented as means ± SEM (n=4; Student’s t-test; p<0.05 is considered significant). **C)** Representative whole mount submucosal preparation from naïve and DSS-treated mouse after in vivo labeling with BMV157 (cathepsin S activity – magenta) and immunostaining with anti-CD68 (green). Note: these samples were collected from Australian mice.

We next analyzed in vivo probe binding in colon tissue, luminal fluids and fecal pellets by in-gel fluorescence (**Fig 3A-F**). In colon tissue, there was a subtle but significant increase in BMV157-labeled cathepsin S (1.6-fold), whereas in fecal pellets and luminal fluids, the increase was more dramatic (5-and 7.8-fold, respectively) (**Fig 3A-F**). Similarly, BMV109-labeled cathepsin S was increased in DSS-treated fecal pellets and luminal fluids by 3.2-and 10.3-fold, respectively, while it was not statistically different to control in proximal colon tissue (**Fig 3A-F**). Cathepsin X, B and L activities were not significantly different between naïve and inflamed colon tissue (**Fig 3A,G**). Immunoprecipitation with cathepsin-specific antibodies was used to confirm the identity of all proteases labeled by BMV109 in colon tissue (**Fig 3H**).

**Figure 3.**
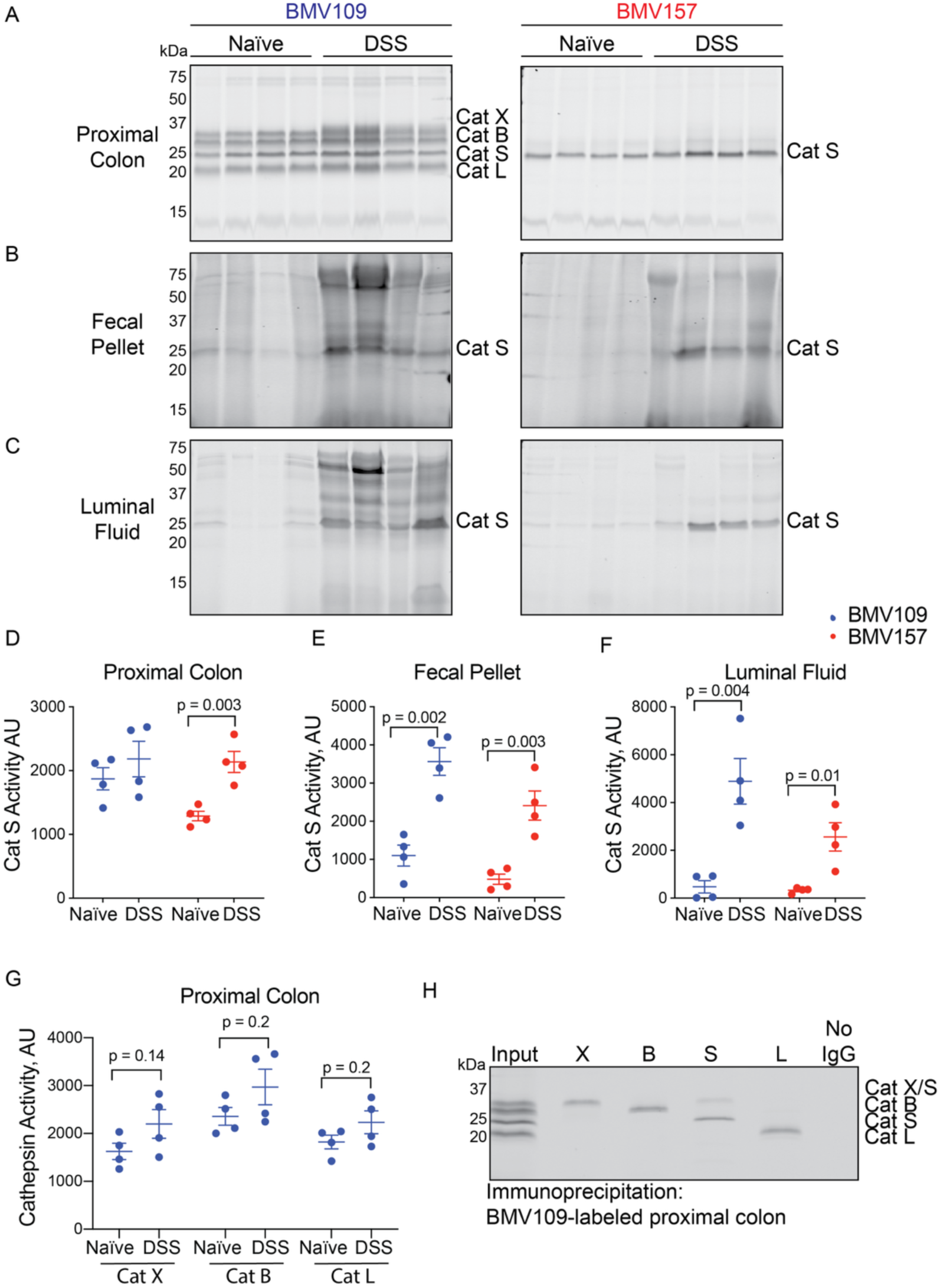
Cathepsin S secretion is increased during DSS-induced colitis. Proximal colons **(A)**, fecal pellets **(B)** and luminal fluids **(C)** from naïve and DSS-treated mice labeled in vivo with BMV109 (left) or BMV157 (right) were analyzed for in-gel fluorescence. Densitometry of cathepsin S labeling by BMV109 (blue) and BMV157 (red) in proximal colon **(D)**, fecal pellets **(E)**, and luminal fluid **(F)**. **G)** Densitometry of cathepsin X, cathepsin B, and cathepsin L labeling in colons by BMV109. Data are represented as means ± SEM (n=4; Student’s t-test; p<0.05 was considered significant). **H)** Immunoprecipitation of BMV109-labeled proximal colon lysates with antibodies for cathepsin X, B, S, and L. No IgG indicates a negative control in which no antibody was added. Note: these samples were collected from Australian mice.

To exclude the possibility that probe uptake was higher in inflamed colons due to factors such as increased blood flow, we also labeled colon tissue lysates, fecal homogenates, and luminal fluids with BMV109 and BMV157 ex vivo. Cathepsin S activity was significantly increased in all sample types with both probes (**Fig 4A-F**). We confirmed the identity of ex vivo-labeled cathepsin S by immunoprecipitation (**Fig 4H**). We also demonstrated that cathepsin S labeling could be abolished by pre-treating the samples with MDV-590 ^22^, a cathepsin-S specific inhibitor (**Fig 4I**). Fecal cathepsin S activity remained elevated after mice were permitted to drink normal water for two days after cessation of DSS treatment (**Fig 4J-K**). By ex vivo labeling, the activities of cathepsins X, B, and L were also significantly increased in inflamed colons (1.7-, 1.3-, and 2.8-fold, respectively; **Fig 4A,G**). By immunoblot, we observed increased levels of pro-cathepsin X in inflamed colons (**Fig 4L-M**).

**Figure 4.**
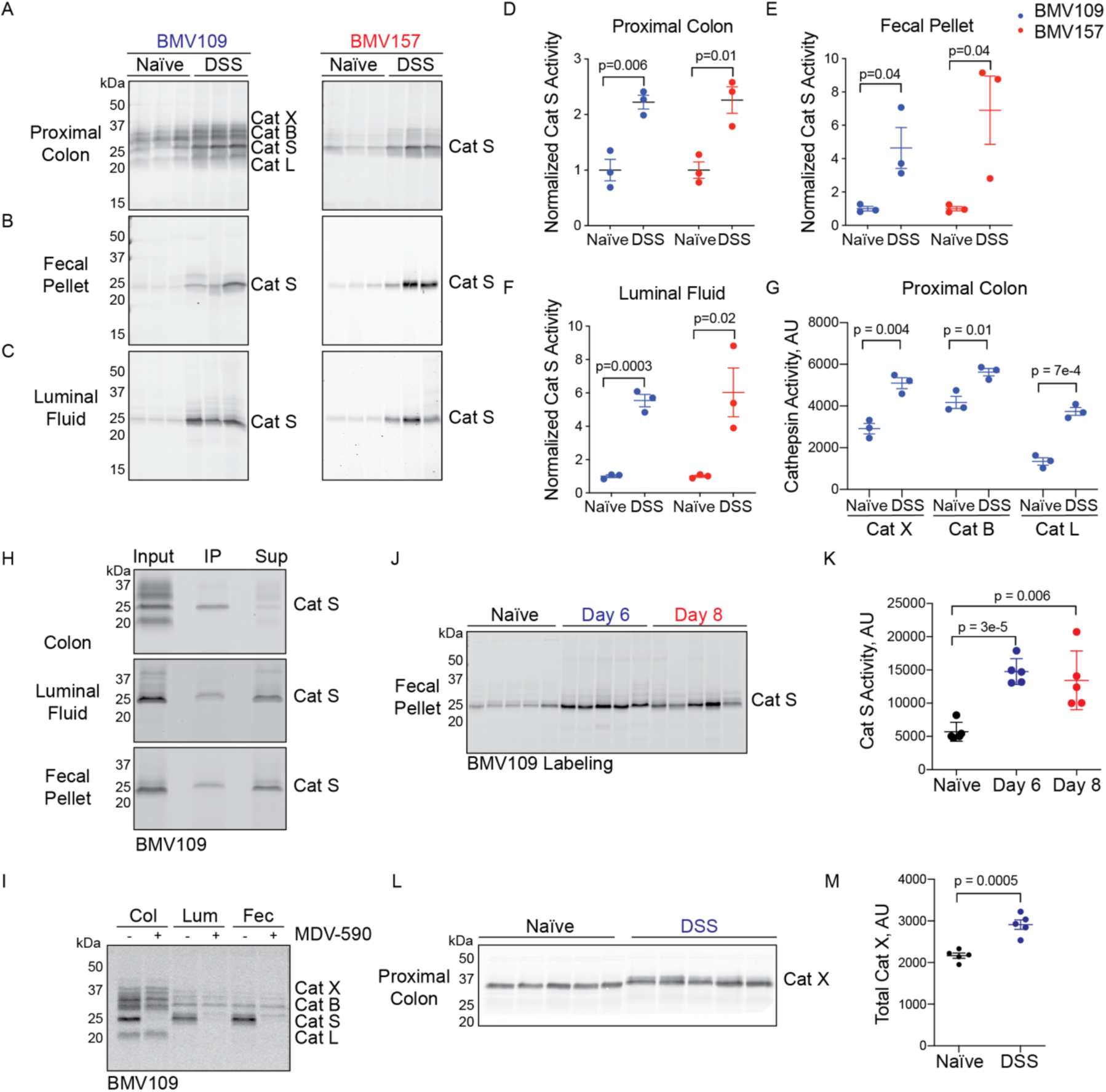
Cathepsin S secretion is increased during DSS-induced colitis. Lysates of proximal colons **(A)**, fecal pellets **(B)** and luminal fluids **(C)** from naïve and DSS-treated mice were labeled in vitro with BMV109 (left) or BMV157 (right) and analyzed for in-gel fluorescence. Densitometry of cathepsin S labeling by BMV109 (blue) and BMV157 (red) in proximal colon **(D)**, fecal pellets **(E)**, and luminal fluid **(F)**. **G)** Densitometry of cathepsin X, cathepsin B, and cathepsin L labeling in (A). Data are represented as means ± SEM (n = 3). **H)** Immunoprecipitation of BMV109-labeled proximal colon lysates, luminal fluids and fecal supernatant with a cathepsin S-specific antibody. IP = immunoprecipitation, sup = supernatant. **I)** Colon lysate (col), luminal fluid (lum) or fecal supernatant (fec) were pre-treated with MDV-590 or vehicle followed by labeling with BMV109 to assess residual cathepsin activity by in-gel fluorescence. **J)** Mice were treated with DSS for 6 days and harvested immediately or after two additional days of normal drinking water. Fecal samples were labeled with BMV109 to measure cathepsin S activity by in-gel fluorescence and the labeling was quantified by densitometry in **(K)**. Data are represented as means ± SEM (n = 5). **L)** Proximal colon tissues from naïve or DSS-treated mice were lysed and then analyzed by immunoblot with a cathepsin X-specific antibody. **M)** Bands were quantified by densitometry. Data are represented as means ± SEM (n = 5). Note: these samples were collected from Australian mice.

To further investigate cathepsin S secretion, we incubated segments of proximal colon from naïve and DSS-treated mice in culture overnight and analyzed the conditioned media for cathepsin S activity. Both groups clearly secreted cathepsin S, although the difference in activity was not significantly different (**Fig 5A-B**). As another potential source of cathepsin S, we also examined mesenteric lymph nodes. While cathepsin S could clearly be detected in mouse lymph node lysates, there was no difference in cathepsin S activity between naïve and inflamed nodes (**Fig 5C**). We also examined cathepsin S activity throughout the GI tract in naïve mice (**Fig 5D**). Cathepsin S activity was much higher in the small intestine and distal colon than the proximal colon. While we have not yet examined small intestine during DSS colitis, secretions from these regions may contribute to the elevated cathepsin S in luminal fluid and fecal pellets. In support of this hypothesis, in rats, DSS has been previously demonstrated to promote inflammation in the small intestine^29^.

**Figure 5.**
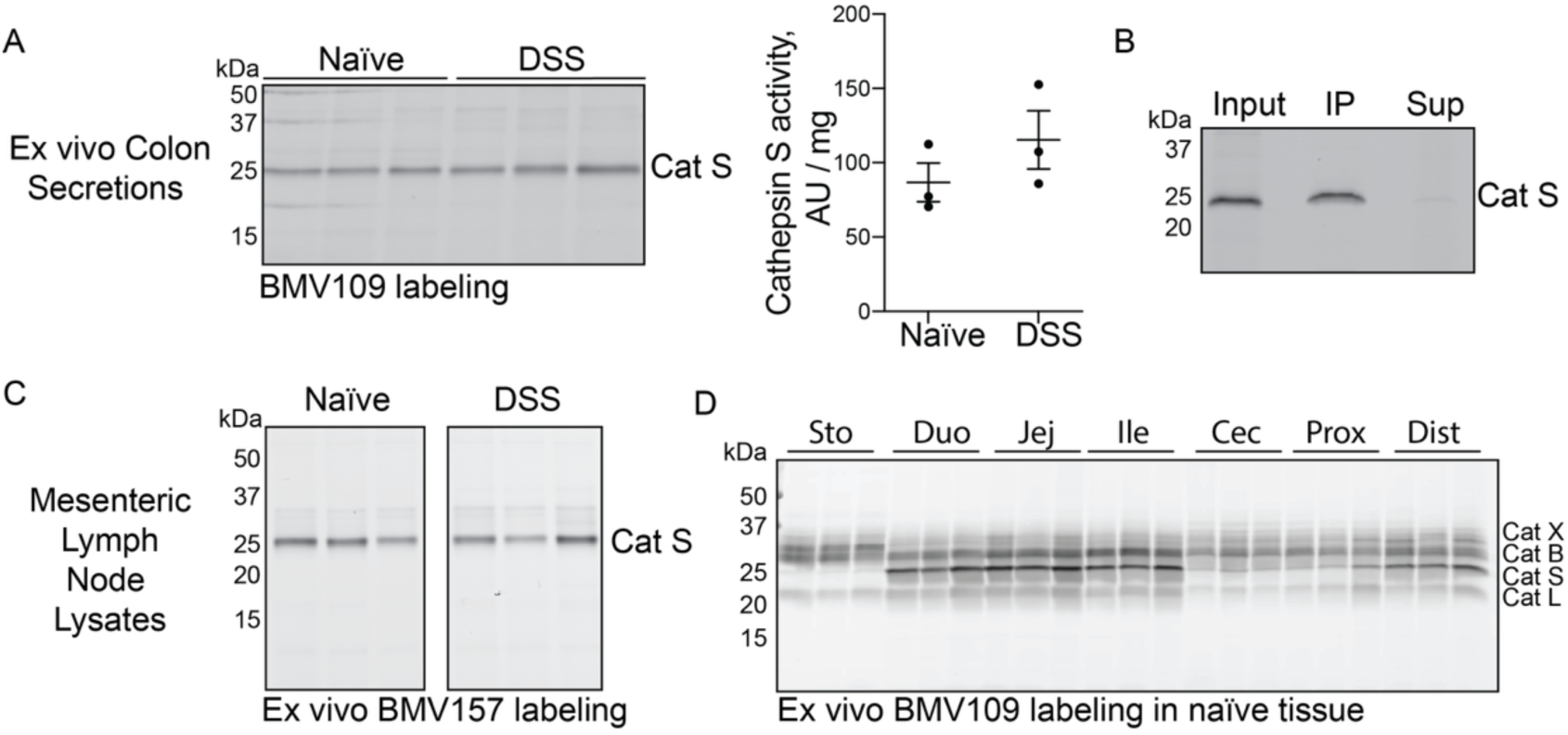
Cathepsin S is secreted from colon tissue. **A)** Proximal colon tissues from naïve or DSS-treated mice were incubated in media overnight. Supernatants were collected, labeled with BMV109 and analyzed by in-gel fluorescence. Labeling was quantified by densitometry and normalized to the weight of each tissue. Data are represented as means ± SEM (n=3). **B)** Supernatants from A were subject to immunoprecipitation with a cathepsin S-specific antibody. IP = immunoprecipitation; Sup = supernatant. **C)** Mesenteric lymph nodes from naïve or DSS-treated mice were lysed, labeled with BMV109 and analyzed by in-gel fluorescence (n=3). **D)** Tissues from the GI tract of naïve mice were labeled with BMV109 ex vivo and analyzed by in-gel fluorescence. Sto = stomach; Duo = duodenum; Jej = jejunum; Ile = ileum; cec = cecum; prox = proximal colon; dist = distal colon. Note: these samples were collected from Australian mice

Collectively, these data demonstrate that cathepsin S is secreted from naïve mouse colons, and this secretion is elevated during acute experimental colitis. The activity of cathepsin S, X, B, and L are also modestly elevated in proximal colon tissue during colitis.

### Cathepsin S deletion, but not cathepsin X, improves symptoms of experimental colitis

We next sought to investigate the contributions of cathepsin S and cathepsin X to DSS-induced colitis using cathepsin-deficient mice. As mice from a Canadian colony were used in these experiments, while Australian mice were used for all other experiments, we first analyzed cathepsin activation in colon tissue, fecal pellets and luminal fluids after DSS treatment. In colon tissues labeled ex vivo with BMV109, activities of cathepsin S, X, and L were significantly increased after DSS treatment in wild-type mice, while cathepsin B activity was unchanged (**Fig 6A, D-G**). In cathepsin S-deficient colons, we did not observe any compensatory changes to the other cathepsins. In the absence of cathepsin X, however, cathepsin L activity was significantly upregulated (**Fig 6A, E**).

**Figure 6.**
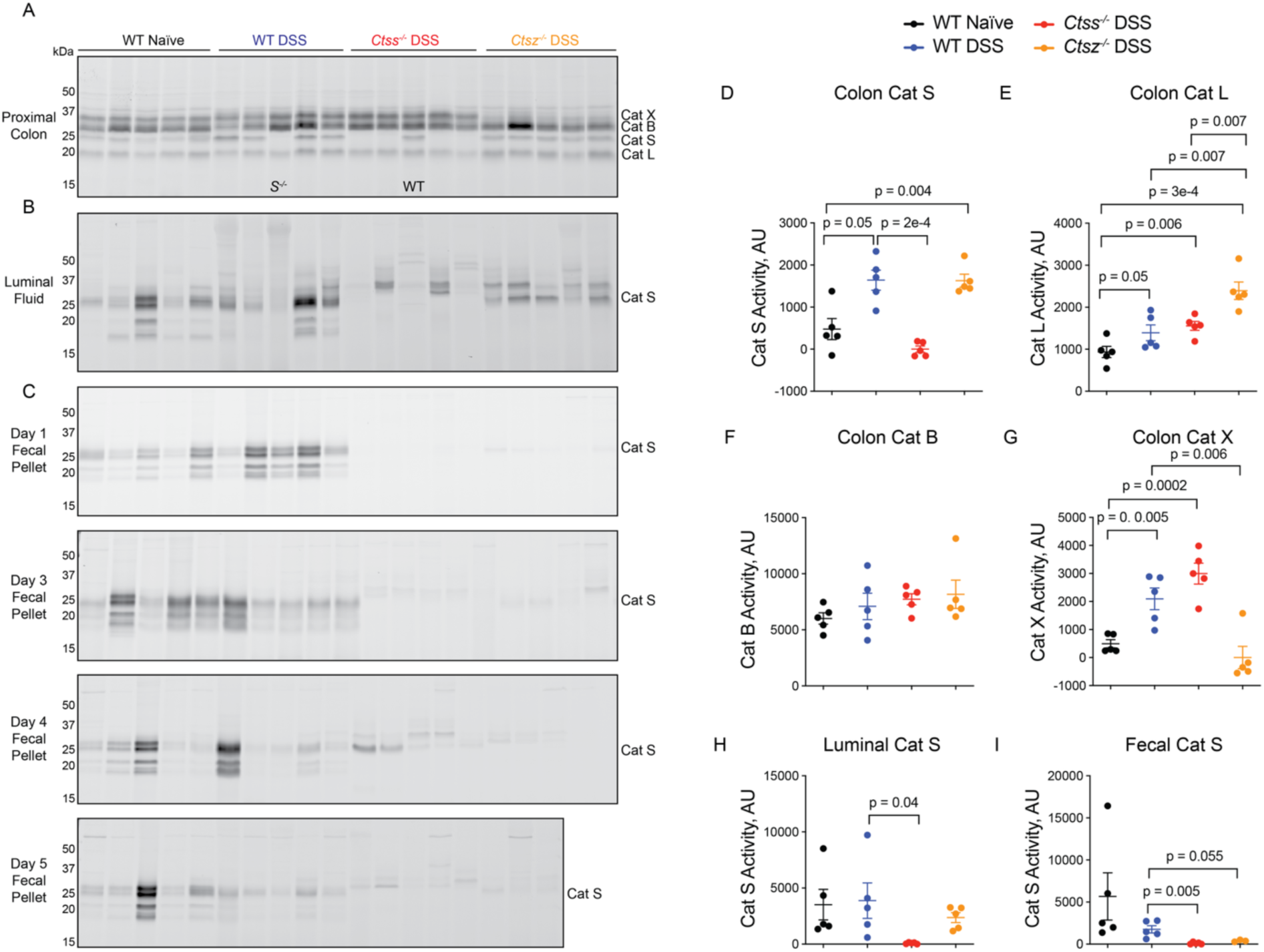
Cathepsin activation in wild-type and cathepsin S-or cathepsin X-deficient mice during experimental colitis. Proximal colons **(A)**, luminal fluids **(B)** or fecal pellets collected at day 1, 3, 4, or 5 **(C)** from wild-type or cathepsin S-or cathepsin X-deficient mice (naïve or DSS-treated) were labeled ex vivo with BMV109 followed by analysis of in-gel fluorescence. Labeling of cathepsin S, X, L, and B in colon tissue lysates was quantified by densitometry (**D-G**), as well as cathepsin S activity in luminal fluids **(H)** and fecal pellets **(I)**. Data are represented as means ± SEM (n = 5). Student’s t-test was used to compare two groups, and p<0.05 was considered significant. Note: these samples were collected from Canadian mice.

Surprisingly, naïve wild-type C57Bl/6J mice from the Canadian colony exhibited much higher protease activity in luminal fluids and fecal pellets than the Australian mice, with a greater diversity of proteins labeled by BMV109 (**Fig 6B-C**). Secretion of cathepsin S did not increase in response to DSS treatment, contrary to what we reproducibly observed in the Australian colony (**Fig 6B-C,H-I**). Moreover, mice deficient in cathepsin S or X had much less fecal cysteine protease activity than the wild-type mice (**Fig 6C,H-I**).

As the changes to protease activity in colon tissue were similar in both colonies, we continued with a comparative analysis of colitis symptoms in wild-type and cathepsin-deficient mice from the Canadian colony. DSS-treated mice lost weight to the same extent, regardless of their genotype (**Fig 7A**). Fecal consistency was likewise similar in all mice, indicating that cathepsin deficiency did not prevent diarrhea (**Fig 7B**). Cathepsin S-deficient mice, however, exhibited less rectal bleeding than wild-type mice or cathepsin X-deficient mice (**Fig 7C,E**). Colon shortening (**Fig 7D-E**) and myeloperoxidase (MPO) activity (**Fig 7F**) were not affected by cathepsin deficiency. Unlike wild-type and cathepsin X-deficient mice, spleens from cathepsin S-deficient mice were not enlarged after DSS treatment (**Fig 7G**). Cathepsin S activity in spleens, however, was minimal and was not altered by DSS treatment (**Fig 7H**). Histological evaluation of colon sections revealed less immune infiltrate, cavitated goblet cells, and crypt disorganization in cathepsin S-deficient mice compared to wild-type or cathepsin X-deficient mice (**Fig 7I-M**).

**Figure 7.**
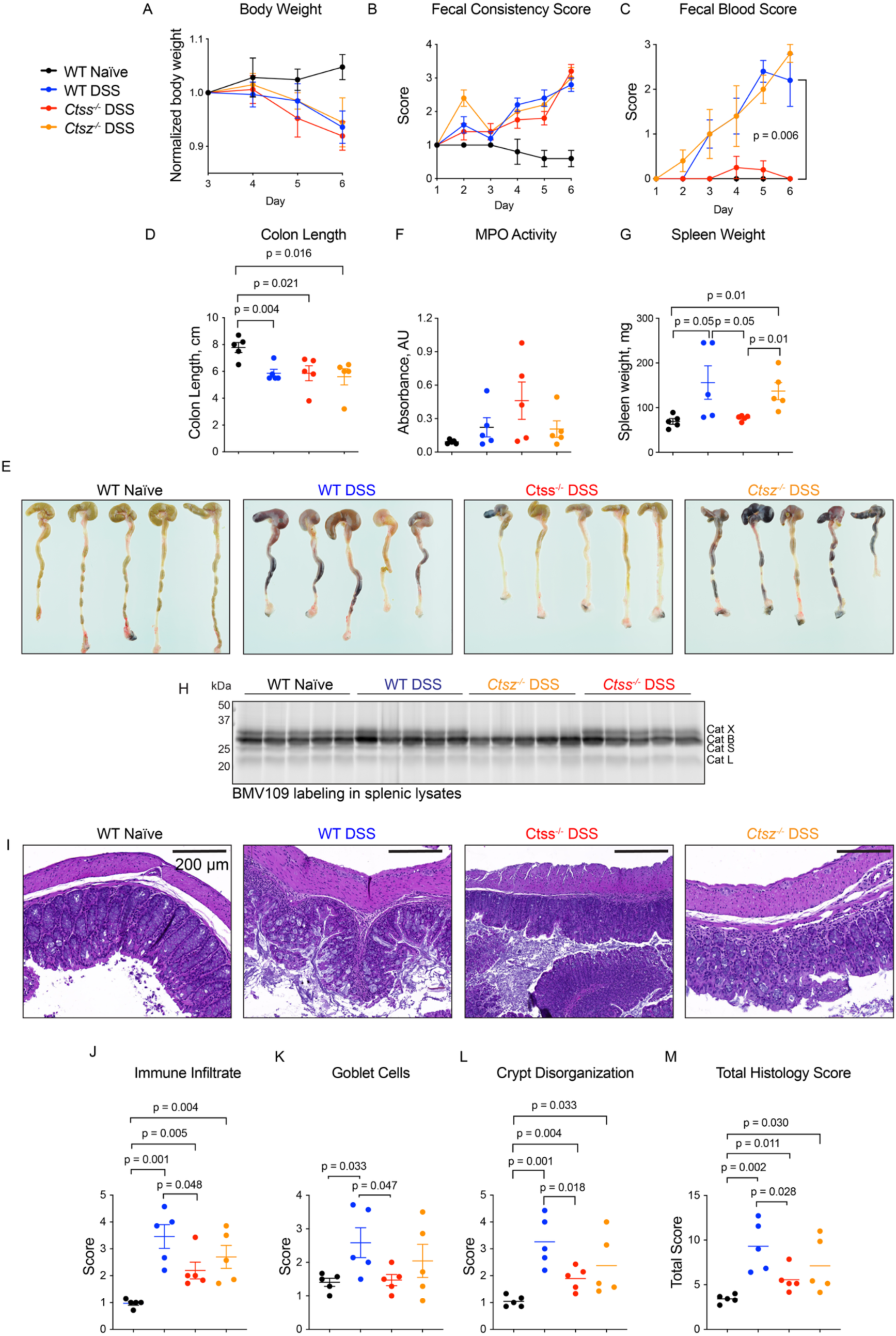
Cathepsin S-deficient mice have improved symptoms of colitis compared to wild-type mice. Naïve mice or mice treated with DSS (wild-type, cathepsin S-or cathepsin X-deficient) were monitored daily for symptoms of colitis, including weight loss **(A)**, fecal consistency **(B)** and fecal blood **(C)**. At end point, colons were excised, measured **(D)** photographed **(E)**, and assayed for myeloperoxidase (MPO) activity **(F)**. **G)** Spleens were weighed at endpoint. Data are represented as means ± SEM (n=5). **H)** Spleens from the same mice were lysed, labeled ex vivo with BMV109, and analyzed by in-gel fluorescence. Note: a portion of this gel was previously published to verify specificity of BMV109 ^20^. **I)** Hematoxylin and eosin staining of colons from naïve or DSS-treated mice (wild-type or cathepsin S-or cathepsin X-deficient). Scoring the sections in (I) for immune infiltrate **(J)**, goblet cells **(K)** or crypt disorganization **(L)** and their combination **(M)**. Data are represented as means ± SEM (n=5). Student’s t-test was used to compare two groups and p<0.05 was considered significant. Note: these samples were collected from Canadian mice.

Collectively, these data suggest that mice from different colonies exhibit variable levels of fecal protease activity at baseline, and differential secretion of cathepsin S in response to colitis induction by DSS. In the Canadian cohort, where mucosal cathepsin S was marginally increased after DSS treatment compared to naïve colons, with no demonstrable increase of cathepsin S secretion, genetic deletion of cathepsin S resulted in marginal improvement of some, but not all, symptoms of colitis. Loss of cathepsin X had no obvious effects in this model.

### Pharmacological inhibition of cathepsin S results in exacerbated colitis

Having observed that genetic deletion of cathepsin S resulted in reduction of some colitis symptoms in the Canadian cohort, we aimed to test the effect of cathepsin S inhibition in the Australian cohort, which exhibited reproducibly enhanced luminal secretion of cathepsin S in response to DSS. We selected the reversible inhibitor LY3000328 as it has previously shown efficacy in a T cell-induced model of colitis, as well as a model of abdominal aortic aneurysm^13,16^. We used a milder model of colitis (2% DSS) in hopes of exacerbating differences in symptoms between treated and untreated mice. Mice were treated daily with vehicle or LY3000328 (30 mg/kg) and symptoms were monitored over time. In naïve mice, LY3000328 had no effect on any of the parameters measured compared to vehicle-treated mice. LY3000328-treated mice with DSS-induced colitis lost significantly less weight than the vehicle-treated controls (**Fig 8A**). No significant differences were observed in fecal consistency (**Fig 8B**) or fecal blood (**Fig 8C**), although treated mice trended towards worse scores at end point. Colon length (**Fig 8D**) and colon MPO activity (**Fig 8E**) were unaffected by LY3000328. LY3000328-treated mice exhibited greater splenomegaly than vehicle-treated controls (**Fig 8F**). Histological evaluation indicated that treated mice had a higher degree of inflammatory infiltrates, cavitated goblet cells, and crypt disorganization than controls (**Fig 8G-K**).

**Figure 8.**
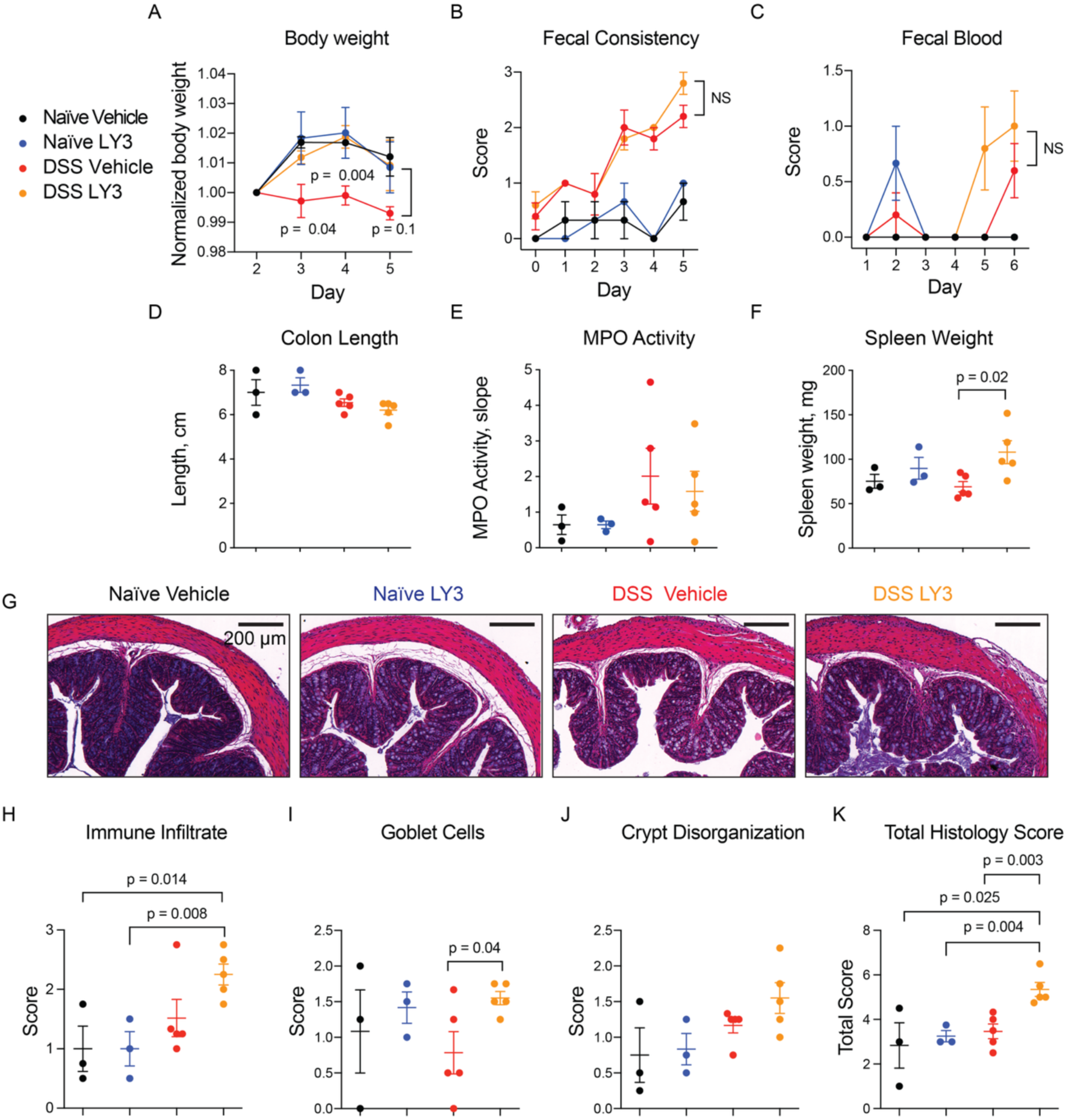
Pharmacological inhibition of cathepsin S exacerbates DSS-induced colitis. Naïve or DSS-treated mice were administered either vehicle or LY3000328 daily over the course of colitis induction. Mice were monitored daily for symptoms of colitis, including weight loss **(A)**, fecal consistency **(B)** and fecal blood **(C)**. At end point, colons were excised, measured **(D)** and assayed for myeloperoxidase (MPO) activity **(E)**. **F)** Spleens were weighed at endpoint. Data are represented as means ± SEM (n=3-5). **G)** Hematoxylin and eosin staining of colons, which were scored for immune infiltrate **(H)**, goblet cells **(I)** crypt disorganization **(J)** or their combination **(K)**. Data are represented as means ± SEM (n = 3-5). Student’s t-test was used to compare two groups and p<0.05 was considered significant. Note: these samples were collected from Australian mice.

Overall, LY3000328 treatment had a negative impact on colitis symptoms, which was contrary to our hypothesis based on the results in the cathepsin S-deficient mice. To ensure that LY3000328 had effectively inhibited cathepsin S activity, we analyzed colon tissue and fecal pellets from these mice using BMV109 ex vivo (**Fig 9A-E**). LY3000328-treated colon lysates contained increased cathepsin S and L activity compared to the vehicle-treated controls (**Fig 9A,C,D**). This corresponded to an increase in total levels of cathepsin S and L as shown by immunoblot (**Fig 9F-G**). Fecal cathepsin S was unaffected by LY3000328 (**Fig 9B,E**). We verified that the reversible LY3000328 was capable of competing for the irreversible BMV109 probe in colon lysates. After LY3000328 pretreatment, residual cathepsin S labeling by BMV109 was blocked in a concentration-dependent manner (**Fig 9H-I**). While LY3000328 exhibited selectivity for cathepsin S, we also observed inhibition of cathepsin X, B, and L in vitro, though at concentrations much higher than the reported window of selectivity. Given that total cathepsin S and L was increased in response to LY3000238, we hypothesize that the drug stimulated increased expression of both proteases. A similar compensatory response has been shown as a response to inhibition of other lysosomal proteases such as legumain and cathepsin L, and in response to lysosomal substrate overload^26,30^.

**Figure 9.**
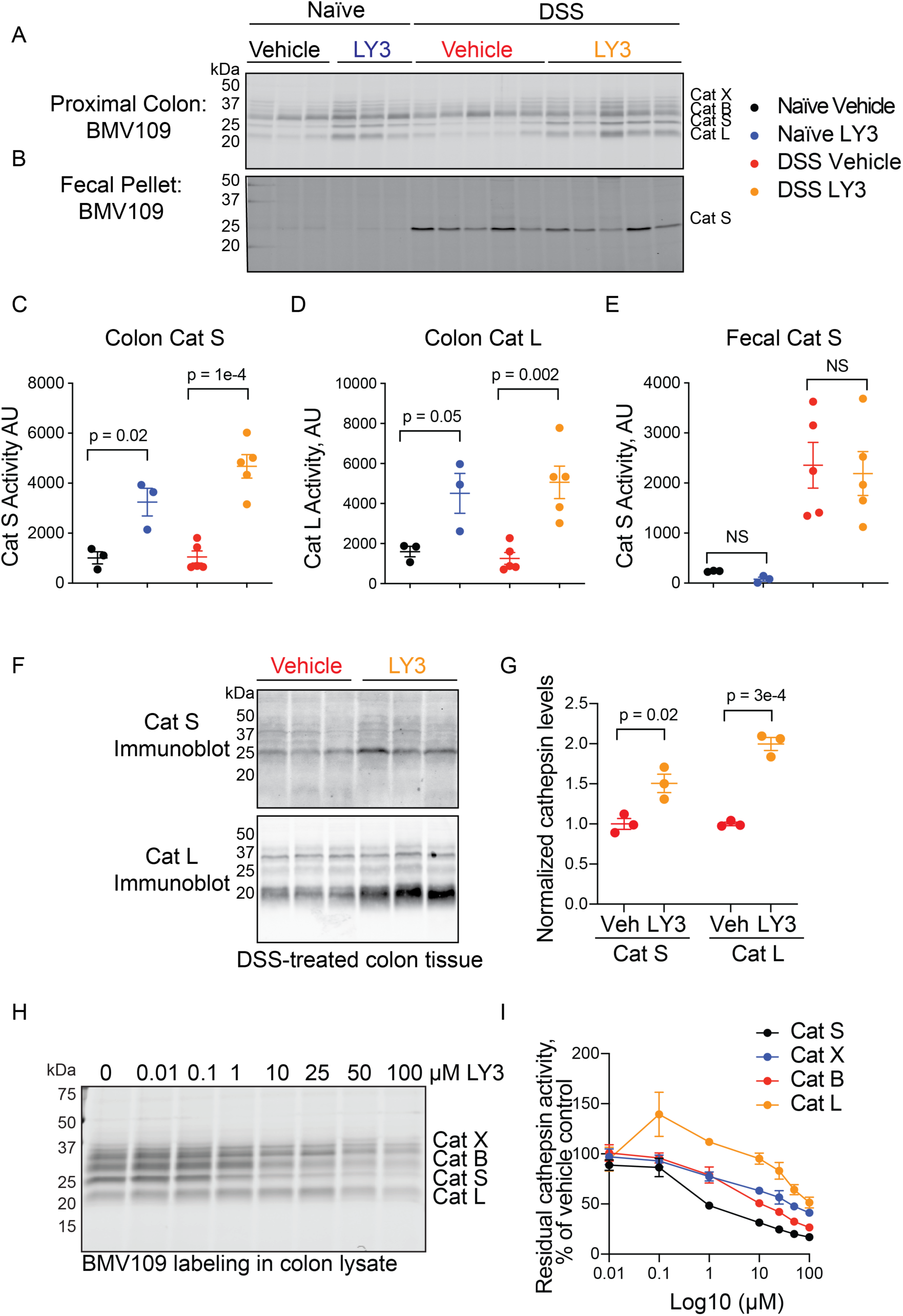
Pharmacological inhibition of cathepsin S increases cathepsin S and L levels. Proximal colons **(A)** and fecal samples **(B)** from mice characterized in Figure 7 were labeled with BMV109 ex vivo and analyzed by in-gel fluorescence. Densitometry of colon cathepsin S labeling **(C)**, colon cathepsin L labeling **(D)** or fecal cathepsin S labeling **(E)**. Data are represented as means ± SEM (n = 5). **(F)** DSS-treated colon tissue lysates from vehicle or LY3000328-treated mice were immunoblotted with antibodies for cathepsin S or cathepsin L and their levels were quantified by densitometry **(G)**. Data are represented as means ± SEM (n=3). **H)** Colon lysates were pre-treated with the indicated concentration of LY3000328 for 30 minutes followed by a 10-minute incubation with BMV109 to detect residual cathepsin activity. Labeling was detected by in-gel fluorescence and analyzed by densitometry **(I)**. Data are represented as means ± SEM (n=3). Student’s t-test was used to compare two groups and p<0.05 was considered significant. Note: these samples were collected from Australian mice.

Collectively, these data revealed an increase in colonic cathepsin S and L activity in response to LY3000328 treatment, and this corresponded to a mild worsening of colitis symptoms. It should be noted, however, that naïve mice did not develop colitis symptoms after LY3000328 treatment, suggesting that increased cathepsin activity alone is not sufficient to drive colitis. While there may be some value in inhibiting cathepsin S as a therapeutic strategy for colitis, it is likely that covalent inhibitors will be required to sustain inhibition as new proteases are synthesized as a compensatory response.

## Discussion

In this study, we employed broad-spectrum and specific activity-based probes to profile cathepsin activity during murine colitis. We observed an increase in cathepsin S activity in luminal fluids and fecal samples in DSS-induced mouse colitis. This corroborates the previous observation that cathepsin S secretion was increased in piroxicam-induced colitis^6^. We demonstrated that cathepsin S can be directly secreted from mouse colon tissue, and that CD68^+^ macrophages are the richest source of cathepsin S.

While enhanced cathepsin S secretion in DSS-treated mice was extremely reproducible in the Australian colony of mice, it was not observed in the Canadian mouse colony. Furthermore, the Canadian wild-type mice exhibited much broader diversity in BMV109-labeled fecal proteases at baseline compared to the Australian mice, while fecal samples from both cathepsin S-and cathepsin X-deficient Canadian mice had little protease labeling at any time point examined. These differences may be attributed to divergence in mouse strains over time or to differences in the gastrointestinal flora populated by the two groups of mice. While the BMV109-labeled proteases in the Canadian wild-type mice were not further characterized, it is possible that some species could be microbially derived.

Two complementary approaches were employed to assess the contribution of cathepsin S to DSS-induced colitis. Genetic deletion of cathepsin S resulted in less rectal bleeding, less splenomegaly, and marginally improved histological scores. Pharmacological inhibition of cathepsin S activity ultimately resulted in exacerbated levels of cathepsin S and L, greater splenomegaly and worsened histological scores. Collectively, these data suggest a role for cathepsin S, and possibly cathepsin L, in promoting colitis symptoms.

Splenomegaly was among the most striking changes upon cathepsin S loss or hyper-activation. This was also observed in a mouse model of periodontitis involving systemic exposure to lipopolysaccharide from *Porphyromonas gingivalis* (PgLPS)^31^. Like colitis, PgLPS exposure leads to splenomegaly in wild-type mice, and this was clearly suppressed in cathepsin S-deficient mice. Cathepsin S deficiency was also marked by reduced CD11c^+^ dendritic cells in the spleen, lower production of IL-6 and less differentiation of Th17 cells. PgLPS was demonstrated to provoke cathepsin S-dependent activation of PAR_2_, which couples to the PI3K/Akt pathway to promote IL-6-dependent splenic Th17 cell differentiation. Another study demonstrated that in corneal epithelial cells, cathepsin S provoked PAR_2_-dependent release of IL-6, IL-8, TNFα, IL-1β, and matrix metalloproteinase 9 (MMP-9)^32^, all of which are important in IBD pathogenesis. In particular, IL-6 signaling is central to the development of intestinal inflammation and its neutralization has been under investigation as a therapeutic strategy for IBDs^33^. IL-6 is also known to synergize with IL-4 to stimulate secretion of cathepsin S from bone-marrow derived macrophages through STAT3/6-dependent mechanisms^23^. This may result in a positive feedback loop to augment cathepsin S secretion during colitis. Furthermore, cathepsin S was also recently shown to cleave and activate IL-36γ, another cytokine with known impacts on antimicrobial defense and barrier function^34^.

In phase I clinical trials, a single dose of LY3000328 exhibited a bi-phasic effect on plasma cathepsin S activity^35^. After a decrease in activity over the first 12 hours, cathepsin S activity returned to baseline followed by an increase in activity that was sustained for at least 48 h. We observed a similar upregulation of cathepsin S activity in mouse colon tissue after daily doses of LY3000328 for five days (**Fig 9A**). Cathepsin L activity was also similarly increased by LY3000328. In vitro, we demonstrated that LY3000328 could compete for labelling by the covalent ABP BMV109. Though it was most potent for cathepsin S, it also inhibited the activity of cathepsin X, B, and L at higher concentrations. Increased protease levels may be a compensatory response to the loss of cathepsin activity, although transcriptional activity of cathepsin S and L was not assessed in this study or in the human trial. Inhibition of another lysosomal cysteine protease called legumain results in transcriptional upregulation of itself and other lysosomal hydrolases, likely through a STAT3-mediated mechanism^30^. We previously showed that legumain accumulates in colon, pancreas and kidney tissue after administration of the small-molecule inhibitor, LI-1^21,36^. As LI-1 is an irreversible inhibitor, however, legumain activity was completely blocked despite the increase in total legumain protein. These data suggest that an irreversible inhibitor might be more appropriate for ensuring sustained inhibition of cathepsin S in vivo and improved therapeutic outcomes.

Therapeutic application of elafin, an endogenous protease inhibitor, has recently been shown to reverse fibrosis-induced intestinal strictures by blocking the ability of cathepsin S to activate PAR_2_, thereby reducing collagen expression^37^. Likewise, legumain inhibition improved fibrosis associated with chronic pancreatitis, but had no clear effects in acute pancreatis or colitis^21,36,38^. We hypothesize that more pronounced effects of cathepsin S blockade made be observed in chronic settings.

We also investigated the contribution of cathepsin X (also known as cathepsin Z) to colitis. This had not previously been explored, and there is limited information on its role in other inflammatory diseases. In the GI tract of naïve mice, cathepsin X activity is highest in the stomach (**Fig 5D**) and its activity may contribute to chronic mucosal inflammation associated with *H. pylori*-induced gastritis and gastric cancer^39,40^. In multiple sclerosis, cathepsin X-deficient mice exhibited less neuroinflammation and secretion of inflammatory cytokines. In a mouse model of silicosis, loss of cathepsin X was associated with reduced inflammation and improved histopathology scores^41^. In this setting, its actions are hypothesized to be extracellular via binding of the RGD integrin-binding motif in its prodomain, as opposed to its catalytic activity. Based on these studies and our observation that pro-cathepsin X expression was increased during colitis, we hypothesized that cathepsin X deficiency would similarly ameliorate symptoms of colitis. On the contrary, cathepsin X knockout mice were statistically indistinguishable from wild-type mice for all symptoms examined. Thus, cathepsin X is unlikely to be a suitable therapeutic target for acute colitis.

## Conclusions

Fecal cathepsin S is increased in patients with UC and CD compared to healthy volunteers. Cathepsin S is secreted during murine colitis, though this observation varied by mouse colony. Fecal cathepsin S therefore warrants further consideration as a diagnostic marker for intestinal inflammation. Genetic deletion of cathepsin S, but not cathepsin X, marginally improved symptoms of colitis, while pharmacological inhibition of cathepsin S by the reversible drug LY3000328 ultimately led to increased cathepsin S and L activity and exacerbated colitis symptoms. Together these data suggest a pathological role for cathepsin S in acute colitis, either through actions upon PAR_2_ or other substrates. Improved inhibitors that can promote sustained and specific inhibition of cathepsin S may have value as a therapeutic strategy for intestinal inflammation as well as visceral pain.

## Acknowledgements

We thank C. Nowell for maintaining the imaging facilities at the Monash Institute of Pharmaceutical Sciences. We thank L. Leone and T. Cardamone from the Melbourne Histology Platform and Australian Phenomics Network for histological services. We thank T. Reinheckel for use of the cathepsin X-deficient mice. We thank R. Kochappan, J. Di Cello, and P. Rajasekhar for technical assistance. We thank E. Lindström from Medivir AB for providing MDV-590. LEM was supported by the National Health and Medical Research Council of Australia (NHMRC, GNT2011119) and a University of Melbourne EMCRA seed grant. NWB was supported by grants from the National Institutes of Health (R01NS102722, R01DK118971, R01DE026806, R01DE029951, RM1DE033491) and the United States Department of Defense (W81XWH1810431, W81XWH2210239).

## Declarations of Interest

NWB is a founding scientist of Endosome Therapeutics Inc. Research in NWB’s laboratory is supported in part by Takeda Pharmaceuticals, Inc.

## Author Contributions

LEM conceived the study, planned all experiments, analyzed data, wrote the manuscript, and contributed funding. BMA, RIC, ARZ, HW, BX and SEC executed the experiments and collected data. RMM assisted with data analysis. RMY, DPP and NWB contributed intellectually, advising on experimental design and data interpretation.

## Notes

### Competing Interest Statement

NWB is a founding scientist of Endosome Therapeutics Inc. Research in the NWB laboratory is supported in part by Takeda Pharmaceuticals Inc.

